# Methotrexate-modulated talin-dynamics drives cellular mechanical phenotypes via YAP signaling

**DOI:** 10.1101/2023.04.07.535979

**Authors:** Debojyoti Chowdhury, Sukhamoy Dhabal, Madhu Bhatt, Debashruti Maity, Soham Chakraborty, Keshav Kant Ahuja, Shreyansh Priyadarshi, Shubhasis Haldar

## Abstract

Methotrexate is a well-known antineoplastic drug used to prevent cancer aggravation. Despite being a targeted therapeutic approach, its administration comes with the risk of cancer recurrence, plausibly through its proven off-target effect on focal adhesions. Since FA dynamics is dependent on force transmission through its constituent proteins, including talin, methotrexate might affect the mechanical activity of these proteins. Here we have combined single-molecule studies, computational dynamics, cell-based assays, and genomic analysis to unveil the focal adhesion-regulating role of methotrexate central to its effect on talin dynamics and downstream pathways. Interestingly, our single-molecule force spectroscopic study shows that methotrexate modulates the bimodal force distribution of talin in a concentration-dependent manner. Steered molecular dynamics reveal that methotrexate-talin interactions alter talin mechanical stability exposing their vinculin binding sites. Finally, we found that methotrexate-regulated talin-dynamics remodel cancer cell mechanical phenotypes like cell polarity, adhesion, and migration by regulating talin-vinculin association-mediated YAP signaling. These results further correlate with genomic analysis of methotrexate-treated patients, demonstrating its clinical importance. Taken together, these findings disseminate the effects of methotrexate-modulated mechanosensitivity of adhesion proteins on cellular events.

---

Methotrexate (MTX) is one of the most renowned anti-neoplastic drugs, used in a wide range of concentrations ranging from <1 μM in adult carcinomas to >100 μM in paediatric cancers^1,2^. However, high doses of MTX lead to severe side effects and are associated with cancer recurrences that might lead to chemoresistance^3–5^. Interestingly, recent studies have shown that MTX increases cancer cell migration in a caveolin-dependent manner, affecting the focal adhesion (FA) dynamics^6,7^. Since FA dynamics is majorly dependent on force transmission through its constituent proteins^8,9^, MTX might have some regulatory effect on the mechanical activity of these proteins. Talin is a pivotal FA protein known to possess a force-dependent structure-function relationship that governs FA dynamics^8,10,11^. The direct involvement of MTX in regulating focal adhesion-mediated mechanical responses thus raises the possibility of altered talin function. However, the MTX effect on Talin dynamics under mechanical control remains unexplored.

The biological activation and downstream signaling of talin significantly depend on the mechanical activity of the R3 domain^12^. Mechanical unfolding of this domain relieves talin from its auto-inhibited form aiding vinculin or actin binding^12^. Here, we have used magnetic tweezers (MT) to explore the effect of MTX on a mechanically-stable talin R3 domain with an unaltered binding mechanism and functions, and observed if it has any systemic consequences on cellular responses. This talin R3 domain has been well-characterized as a model FA protein in force-spectroscopic studies due to its role as a primary mechanosensor for talin-based adhesion maturation^10,11,13^. In our MT setup, the talin R3 domain is inserted between N-terminal HaloTag and C-terminal AviTag for their attachment between a glass surface and a paramagnetic bead, respectively^10,11,14,15^. Owing to this magnetic manipulation, the force can be specifically applied on the substrate (Fig. 1A) and has been calibrated using the magnet law which has been extensively described in the literature^14–19^. While applying a force-ramp scan on the talin domain with a loading rate of 3.1 pN/s, we observed their unfolding event as a rapid extension at a certain force, which can be estimated from the force-extension curve. Notably, talin unfolding has been observed to exhibit two different unfolding force populations-a well-known ~20 nanometer (nm) extension at a low force of ~9 pN and a ~30 nm step size at ~31 pN (Fig. 1B). We obtained a contour length of 35.6±0.6 nm and Kuhn length of 0.93±0.01 nm while fitting these different extension values to the freely jointed chain model, suggesting that generation of these different step sizes belong to talin R3 unfolding (Supplementary Fig. 1). Therefore, the bimodal unfolding force being an intrinsic character of structural stability indicates that talin domain exists as two conformers with a difference in their mechanical stability.

**Figure 1:**
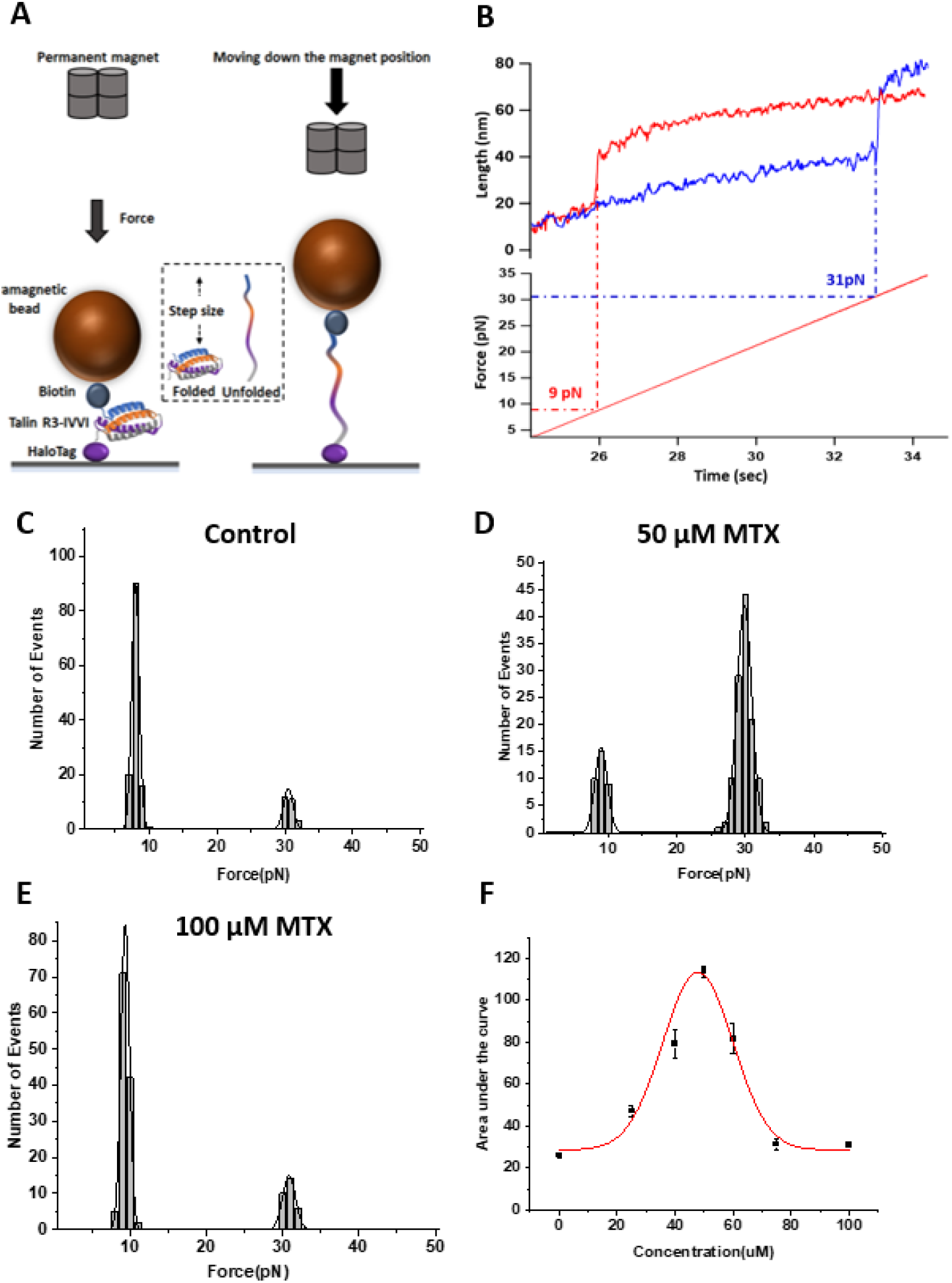
Magnetic tweezers revealed bimodal unfolding force of talin R3: **(A) Instrumentation:** The designed talin R3 construct is tethered between glass surface and paramagnetic bead through the HaloTag and biotin-streptavidin chemistry, respectively. The applied force is estimated as an inverse function of the distance between paramagnetic bead and the permanent magnet. Talin domain remain folded in low force, while become unfolded after applying the high forces; and the difference between these two states is defined as step-size. Image is not scaled. **(B) Mechanical unfolding of talin:** Applying the force at a loading rate of 3.1 pN/s on talin domain results in its unfolding at particular force. Surprisingly, talin exhibits a discrete mechanical strength by unfolding at two different force ranges: one at 9 pN and another at 30 pN. This unfolding pattern indicates the presence of two different mechanical conformers of talin under physiological force, which is also reflected into their extension. **(C to E) Unfolding force histogram of talin with MTX: (C)** Histogram of the estimated talin R3 unfolding forces without MTX has been plotted. The solid black lines represent the fitting with a double Gaussian equation with the data. Unfolding histogram shows two distinct force ranges: first peak indicates the low force population of talin, while the next one corresponds to talin high force population. Approx. 88% of the total population (n=151) are measured in a low force population and the rest of the population in the high force population. This Bimodal unfolding force distribution has been observed to change upon the addition of MTX. **D.** Interestingly, at 50 μM MTX, the higher unfolding force population is approximately 79% of the total population (n=150). **C.** However, further increasing MTX concentration to 75 μM again declines the higher force population with the predominant presence of talin low force population (n=151). **F.** The area of talin high force population is plotted against MTX concentration, showing a biphasic effect on talin mechanical stability.

Given the ability to unfold at two different force regimes, we further studied how the relative population of talin force distribution could be modulated while interacting with MTX under physiological tension. Since talin has been used as a model FA protein in force spectroscopy techniques^8,10,20,21^, we systematically investigated the MTX effect on the talin mechanical stability. To understand the fundamental principle of how MTX could impact talin mechanical stability, we titrated the substrate up to a high MTX concentration, where their effect becomes saturated. Fig. 1C shows that talin R3 mostly unfolds at ~9 pN force as the predominant population with a rare unfolding population at ~31 pN. Surprisingly, MTX tightly regulates this bimodal distribution in a concentration-dependent fashion: promoting optimal unfolding at higher force in the presence of 50 μM MTX (Fig. 1D). However, above this concentration, MTX has been observed to decrease the mechanical stability, almost upto that of talin R3 without the drug (Fig. 1E and 1F), suggesting a prominent biphasic effect of MTX on talin mechanical stability. To crosscheck the specificity of the MTX effect, we further studied a non-FA protein called protein L, which has been extensively studied using force spectroscopy techniques^14–16,19,21^. Notably, we found that MTX do not exhibit any significant effect on protein L mechanical stability, indicating a talin-specific mechano-regulatory activity (Supplementary Fig. 2). Our single-molecule experiments resolved a surprising behaviour of talin mechanical stability with interacting MTX molecules, which revealed a generalized question that how MTX molecules interact with talin domain under force. For a possible explanation, we performed steered molecular dynamic simulation to understand the MTX binding to talin and dynamic nature of their interaction under force similar to our experimental set up.

Talin contains 11 Vinculin binding sites (VBS), two of which (VBS4 and VBS5) is found in the second, and third helix of the R3 domain respectively^22^. The mechanical unfolding of this domain causes vinculin binding, activation, and subsequent vinculin binding to the other VBS^12^. To check the dynamics of these VBSs, that will confer talin’s biological influence, we have performed SMD (NAMD) simulation both in equilibrium and stretched condition at 0, 1, 50, and 100 μM MTX concentration (Fig. 2). Our SMD study with a constant velocity pulling at 0.05Å/ps (0.0001 Å/timestep) suggested that talin R3 unfolds at ~550 pN, while it increases to ~595-768 pN range in the presence of MTX (Fig. 2). Analysis of SMD trajectories indicates that helix 2 and 3 contribute most prominently to the talin mechanical stability and unfolds after helix 1 and 4. Therefore, we speculated that increasing mechanical stability due to MTX would result in the strengthening of helix 2 and 3. Surprisingly, in presence of 1 μM MTX, helix 3 unfolds faster than the control; however, the overall R3 domain gains mechanical stability due to helix 2 and helix 1 stabilization. We found that intramolecular H-bond and intramolecular salt-bridge interactions account for the helix specific stability (Fig. 2D; Supplementary Fig. 3). We performed additional analyses to get a view of talin-MTX interaction dynamics. Indeed, we found that these interactions are not long-term and suggest off-target specificity despite of H-bonding interaction between talin and MTX.

**Figure 2:**
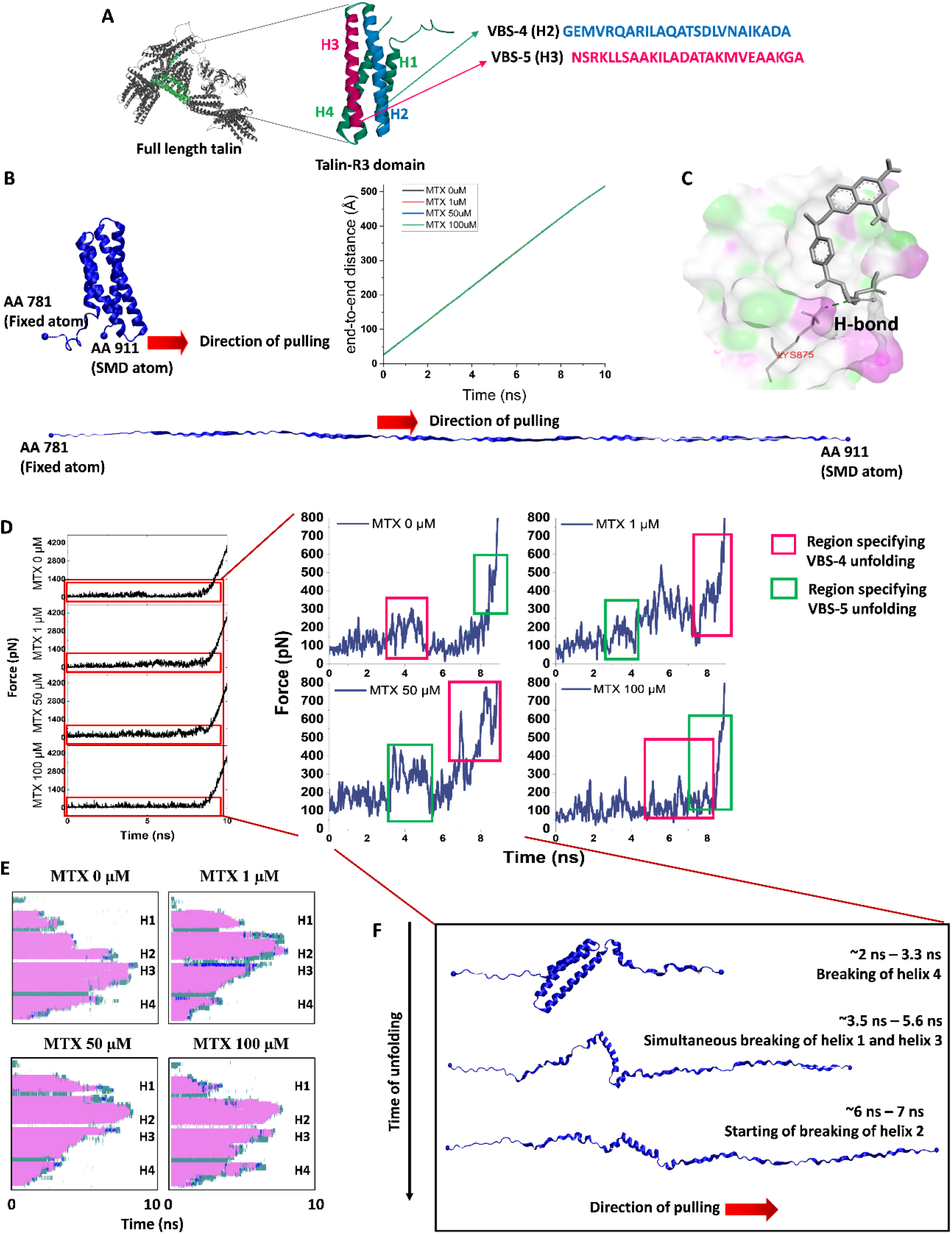
Simulation revealed the difference in talin unfolding force with and without MTX: **(A) Talin structure:** R3 is the initial gatekeeper of all talin domains during force transmission. R3 domain contains 4 helices, among which helix 2 and helix 3 contains the cryptic vinculin binding sites (VBSs). These sites become exposed upon mechanical unfolding of R3 domain. The autoinhibited full length-talin structure is taken from alpha fold. The R3 domain structure is taken from the PDB ID: 2L7A. **(B) Analysis of NAMD simulation trajectory with MTX:** During steered molecular dynamics, the Cα of aa 781 was kept fixed and Cα of aa 911 was pulled with a virtual spring at constant velocity. The end-to-end distance after 10 ns unfolding was found to be ~516 Å. **(C)** Snapshot image from talin-MTX simulation showing stable H-bonding between hydrophilic amino acid residues (lysine here) of talin and charged groups of methotrexates. **(D)** Force (pN) vs time (ns) graph for SMD simulations of control and talin in different drug concentration. The major unfolding events are represented in the zoomed section. Regions corresponding to the unfolding of two VBS are shown in colored boxes accordingly. In 1, and 50 μM conc. the 5^th^ VBS opens before the 4^th^ one which is opposite to the unfolding of these VBSs in control and in 100 μM MTX. The supporting files and information are available as source data. **(E) Secondary structure modulated by MTX interaction:** Perturbation of secondary structure in presence or absence of drugs. H1-H4 indicate the four alpha-helices accordingly. **(F) SMD snapshots** to represent the major talin unfolding events in presence of 50 μM MTX.

Talin mechanical stability has a well-characterized function in controlling the cellular responses^23–27^. A major phenotypic property of a mechanically-induced migrating cell is its ability to change shape, which is an indicator of actomyosin-mediated traction force generation ^28^.We have measured circularity index (or cell shape index) as a quantitative parameter for traction-induced deformation in cells^29^. The circularity index is scaled as such: 0 means a completely elongated cell representative of higher mechano-response, while 1 signifies full circularity and no migration-induced directionality. Both HEK293T and HeLa cells have been observed to transform their structure from spherical to more elongated while increasing the MTX concentration upto 50 μM, which has been estimated as decrease in the circularity index (Fig. 3A, 3B). Interestingly, upon further increasing the concentration to 100 μM, these cells again revert back to spherical shape (Fig. 3B). This strongly signifies that at 1 μM and 50 μM MTX, the elongated cells generate higher traction force, and thereby the migration rate. Since mechanical force acting on individual FA proteins could regulate the adhesion dynamics; change in the FA protein mechanical stability could influence the adhesion dynamics as well. To investigate the effect of MTX-modulated talin stabilization on cellular behaviour, we determined the adhesion dynamics and migration rate of cancer cells^23^. We observed that both the cell types exhibit highest cell-matrix adhesion at 50 μM MTX, which surprisingly decreases upon increasing the concentration to 100 μM (Fig. 3C and 3D). Since cellular migration is cell-matrix adhesion-dependent^9,30–32^; we have also quantified the migration rate through scratch-induced migration assay and found that it is strongly correlated with the adhesion: being highest at 50 μM MTX, followed by ~2 folds decrease with 100 μM MTX (Fig.3E, 3F and inset). This is relevant as mechanical strength of talin R3 is well-known to be proportional with the cellular migration and the migration rate in the presence of 100 μM MTX becomes slower due to the change in FA composition and their dynamics^23^. With 50 μM MTX, talin R3 possibly unfolds at higher forces, especially in FA transmitting high-magnitude force. Such unfolding at relatively high force not only exposes cryptic vinculin-binding sites but also unfold other talin domains that could potentially trigger downstream signaling pathways to increase the migration rate.

**Figure 3:**
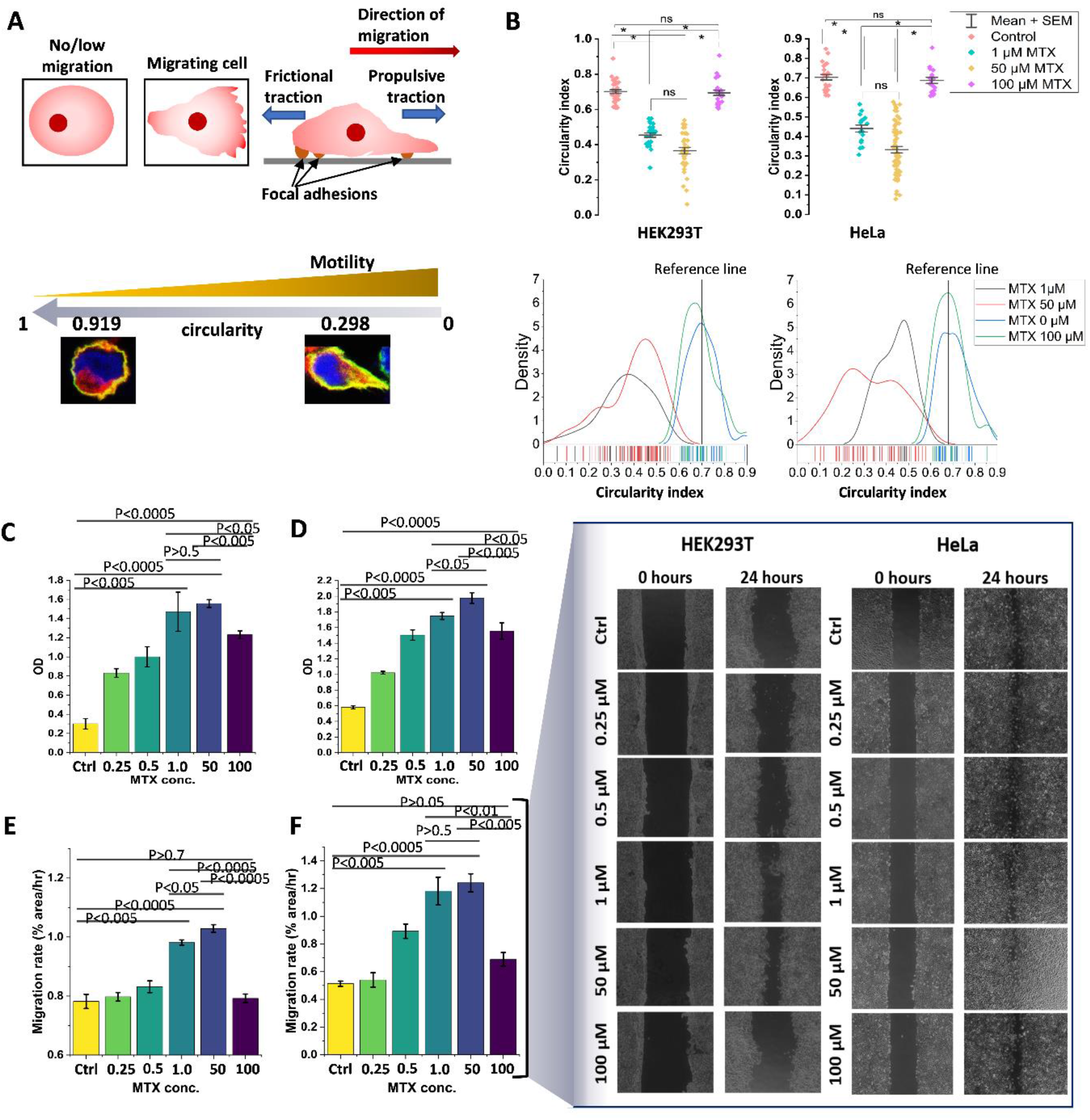
MTX-induced adhesion and migration in cancer cells: **(A and B) Circularity index of migrating cancer cells:** In the absence of MTX, cells are mainly spherical and thus the circularity index is ~1. Upon the treatment with MTX at various concentration, we observed that the circularity index decreases steadily, becoming lowest at 50 μM MTX. However, upon increasing the concentration to 100 μM MTX, the circularity index increases to almost control. Both HEK293T and HeLa cells show similar pattern in the drug-induced circularity index. The reference lines indicate most probable circularity index of both the cell lines in control groups: 0.7 (HEK293T) and 0.64 (HeLa). **(C and D) Adhesion assay:** To check the cell-substrate adhesion, 96 well plates were coated with fibronectin and cell adhesion were treated with different MTX concentrations. Non-adhered cells were washed with PBS after 12 hours and adherent cells were stained with crystal violet to measure OD at 595 nm. Cell adhesion increases gradually from 0 to 50 μM MTX concentration and then decreases at 100 μM MTX. **(E and F) Migration rate:** (Inset magnified) A linear scratch wound was made in a confluent HEK293T and HeLa cell monolayer. Then both the cells were treated with MTX at different concentrations; and incubated in a 37°C CO_2_ incubator for 12 h. For both the cells, wound healing images were taken by bright field microscope and estimated as migration rate (% area/h). **(C and D)** Bar graph shows the estimated cell migration rate in HEK293T and HeLa cells. Data were represented as mean ± S.E.M, n=3. Statistical significance has been given in the graph. (Ctrl: Control).

We co-transfected two different cell lines-HeLa and HEK293T cells with mCherry-vinculin and EGFP-talin for their overexpression; and then treated with MTX at different concentration. Similar to our migration and adhesion studies, we found highest talin-vinculin colocalization at 50 μM MTX due to their strongest association in cells which decreases significantly in control as well as at 100 μM MTX (Fig. 4). The colocalization coefficient is also significantly high in presence of 1 μM MTX plausibly due to talin-vinculin interaction at 5^th^ VBS as indicated by our simulation studies. These data strongly suggest that MTX-modulated talin stabilization could function as a critical regulator of adhesion turnover. To further investigate this, we mapped the size of focal adhesions in talin-GFP and vinculin-mCherry co-expressed cells. Interestingly, in presence of 1 and 50 μM MTX, the focal adhesions formed in both the cell lines comprise two different populations with different focal adhesion size (denoting nascent adhesions and matured adhesions) (Supplementary Fig. 4). The higher size population is absent in control as well as 100 μM MTX-treated group. More importantly, in 1 and 50 μM MTX, the larger sized focal adhesions are increased, which is in agreement with our migration studies that supports the previous finding that with increasing FA size, migration rate also increases^33^. Altogether, in presence of 1-50 μM MTX, the dynamics of focal adhesion maturation and abortion is higher, and thereby producing significant amount of frictional and propulsive force helping the cells to migrate, which is minimal in control and in 100 μM MTX treated cells. In addition to the talin-vinculin accumulation, drug-modulated talin stability also affects cell shape and size. The non-significant difference between the control and 100 μM MTX population in all these mechanical phenotypes suggest that the drug molecules at highest studied concentration almost rescue the untreated cancer cell behaviour.

**Figure 4:**
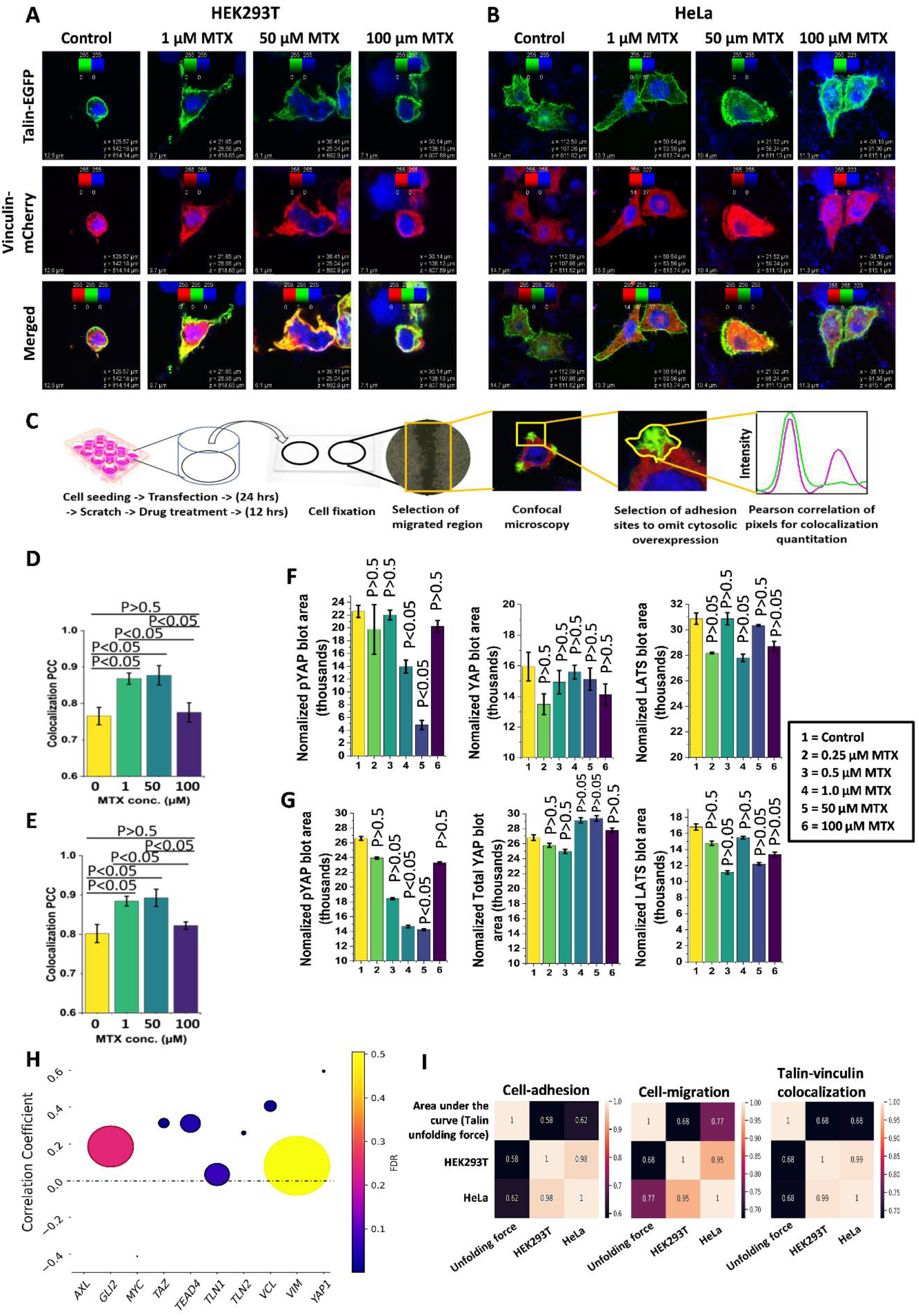
MTX-modulated talin mechanical stability reflects into its functional outcome in cancer cells: **(A and B) Talin-vinculin colocalization:** HEK293T and HeLa cells were co-transfected with mCherry-Vinculin and EGFP-Talin1 to visualize their association in cells. Upon 24 h of transfecting the cells, they were treated with MTX at different concentration; and incubated at 37°C for 12 hrs in a CO_2_ incubator. Cells were fixed with 4% paraformaldehyde, stained with Hoechst dye and mounted for confocal microscopy. **(C)** For quantifying colocalization in the migrating cells, the seeded cells were transfected and incubated for 24 h inducing vertical scratch thereafter. MTX treatment was given immediately after the scratch and further incubated for 12 h. The coverslips from 12 well-plates were taken out and fixed according to the protocol. Only the cells near the migrating ends (scratched zone were focussed for confocal microscopy. Cellular regions denoting focal adhesion assembly (polar tips) were further analyzed for talin-vinculin colocalization using Pearson’s correlation coefficient (pixels). **(D and E)** Captured images were analyzed and colocalization was determined through Coloc2 plugin of Fiji ImageJ software to plot the bar graph. Data were represented as mean ± S.E.M, n=3. **(F, G) Normalized western blot quantitation of pYAP, YAP, and LATS with respect to β-actin (loading reference).** Total YAP protein expression has been observed to be random while increasing the administered MTX concentration from 0.25 upto 100 μM for both HEK293T and HeLa cells. Same no-pattern relative expression was observed for LATS. By contrast, phosphor-YAP expression decreases with the concentration, being minimum at 50 μM. Both the cell lines possess similar pattern in the phospho-YAP expression. Data were represented as mean ± S.E.M, n=3. **(H)** We found a strong positive correlation of talin-vinculin association and YAP expression with MTX administration within the range of 0-50 μM MTX. The samples data were extracted from GDSC and CTRP databases. **(I) Statistical correlation of cellular responses with talin mechanical stability obtained by magnetic tweezers:** Pearson’s correlation coefficient plotted to show the correlation between area under the curve of talin mechanical stability, cell-adhesion, cell migration and talin-vinculin colocalization in both the cell lines.

Previous reports suggest that higher talin-vinculin interactions due to the extracellular mechanical cues can guide nuclear localization of YAP^34,35^ hence, regulating mechanical responses. We speculated that MTX induce structural and mechanical perturbation in talin might regulate YAP activity in a vinculin-dependent manner. Thus, we performed western blot analysis to check YAP activity in different MTX concentration (Supplementary Fig. 5). We found that YAP protein expression is not significantly altered in any of the concentrations (Fig. 4F). On the other hand, YAP phosphorylation decreases gradually upto 50 μM concentration of MTX and becomes almost similar to control at 100 μM MTX (Fig. 4G). The protein expression of LATS, an enzyme responsible for phosphorylation and hence RUNX3-dependent degradation of pYAP^36^, is also non-significantly altered in an MTX concentration-dependent manner. This suggests that MTX concentration-dependent mechanochemical regulation neither impacts YAP expression nor LATS expression-mediated YAP degradation. Accordingly, YAP can only escape phosphorylation by nuclear transport at a rate faster than YAP phosphorylation. We assume that MTX-induced talin mechanochemical perturbations lead to traction force generation that might further help mechanical opening of nuclear pore complex through talin-vinculin-actin-laminin axis helping YAP nuclear localization. Therefore, our study suggests that MTX regulates YAP activity in a concentration-dependent manner which might be due to MTX-driven mechanically regulated talin-vinculin association.

To revisit our hypothesis, we cross-checked the talin-dependent Hippo-YAP pathway through genomic analysis of MTX-treated samples. We considered samples from GDSC^37^ and CTRP^38^ databases and limited our study to samples where 0-50 μM MTX were administered. Indeed, we found a strong positive correlation of talin-vinculin association and YAP expression with MTX administration (Fig. 4H, Supplementary Fig. 6). However, recent studies suggested that YAP can promote both tumor invasion, migration and apoptosis^39,40^. While the former happens in a TEAD-dependent manner leading to the activation and transcription of AXL (a receptor tyrosine kinase), Gli2 (Glioma-Associated Oncogene Family Zinc Finger 2), and VIM (Vimentin), the later occurs due to Myc overexpression. In both the MTX-treated sample cohort, we found a positive correlation of TEAD4 and YAP/TEAD4-dependent transcriptomic products. However, Myc is negatively correlated with a significant FDR, suggesting that MTX treatment upto 50 μM might impede Myc-mediated apoptosis and promote cell migration (Fig. 4H). To further extend our analysis to in-vivo context, we performed transcriptomic analysis with whole genome data of TCGA patients, who underwent methotrexate therapy. Interestingly, this study also suggests that YAP signaling components and their transcriptomic products that are related to increased mechano-response, increase significantly in MTX-treated patients. However, we found a Myc downregulation trend similar to the cell line-based studies indicating a plausibly lower apoptotic signal. Overall, we have observed a strong correlation area under the curve of talin mechanical stability, cell-adhesion, cell migration and talin-vinculin colocalization in both the cell lines (Fig. 4I).

Although it is consensus that only talin R3 unfolding can cause vinculin binding and activation, Atherton et al. has shown that talin-vinculin interaction is force-independent and activated vinculin can even bind to the mechanically stable mutant of R3 irrespective of its unfolding^41^. Moreover, force transmission through talin and its unfolding are found to be independent^42^. Therefore, it is very much possible that vinculin can also bind to the MTX-induced mechanically stable R3 (at 50 μM concentration). To ameliorate the existing models, our single molecule and computational studies combined with cellular and genomic assays indicate that Vinculin can bind the mechanically stable R3 domain in its high-force conformation. This binding is plausibly more dependent on VBS-5 than VBS-4 as we found a strong correlation between VBS-5 unfolding and increased talin-vinculin colocalization even if VBS-4 is gaining more mechanical stability. Further activation of vinculin and subsequent relief of talin-autoinhibition might cause mechanical alterations in cellular architecture saving YAP from LATS-mediated phosphorylation. This plausibly leads to YAP nuclear localization choosing the YAP-TEAD4 axis to promote transcription of cell polarity and migration-aiding genes rather than apoptosis-inducing genes. Due to the predominant α-helical folds in most of the adhesion proteins at cell-substrate adhesion, this finding could have significant insight for reconstructing their mechanical stability as a plausible therapeutic choice for chemotherapeutic drugs.

## Materials and Methods

### Protein expression and purification

Talin construct was made by BamH1, KpnI, and BglII restriction sites in pFN18a expression vector, as described in our previous studies^10,11,17^. The construct was transformed into *Escherichia coli* BL21 star (DE3) cells, followed by growing them in LB media at 37°C and inducing them by 1 mM isopropyl β-D-thiogalactopyranoside (IPTG, Sigma-Aldrich) overnight at 25°C. The cell pellets were resuspended in 50 mM sodium phosphate buffer (pH 7.4) supplemented with 300 mM NaCl and 10% glycerol and lysed by cell homogenization (Constant systems). Then the proteins were purified from the cell lysate through Ni^2+^-NTA affinity column (ÄKTA Pure, Cytiva). Then the His-tagged eluted protein was biotinylated using Avidity biotinylation kit and recommended protocol, followed by size exclusion chromatography with Superdex-200 increase 10/300 GL column, and finally, eluting the protein in Na-phosphate buffer with 150 mM NaCl and 10% glycerol.

### Flow chamber preparation

The cover glasses (bottom) were sonicated with Hellmanex III (1.5%) and washed with double distilled water. Then it was treated with a mixture containing concentrated HCl and CH3OH, followed by concentrated H2SO4. Then the cover glasses were washed with double distilled water, boiled and dried in hot air oven. Cover glasses were then activated by silanization with an ethanol solution supplemented with 1% v/v (3-Aminopropyl) trimethoxysilane (Sigma Aldrich, 281778) for 15 minutes, followed by washing for few times with ethanol to remove unreacted silane from the glass surface. Finally, the cover glasses were dried and baked at 65°C. Similarly, coverslips (top) were sonicated with 1.5% Hellmanex III for 15 minutes, followed by washing with ethanol and baked at 65°C for 10 minutes. Then both the top and bottom glasses were sandwiched, separated by parafilm template between the two glasses. After sandwiching, the chambers were vigorously flushed with glutaraldehyde and kept to react for an hour. A PBS solution containing the reference bead (2.5-2.9 μm, Spherotech, AP-25-10) was passed and then incubated for 15 minutes, followed by vigorous washing with PBS. Then HaloTag(O4) ligand (Promega, P6741) were passed and incubated overnight at room temperature. Finally, to remove any non-specific interaction, chambers were washed with blocking buffer (20 mM Tris-HCl, 150 mM NaCl, 2 mM MgCl_2_, 0.03% NaN_3_, 1% BSA, pH 7.4) for ~5 hours at 25°C.

### Single-molecule magnetic tweezers experiments

Single-molecule experiments were done on custom-made magnetic tweezers built on a Zeiss Axiovert S100 microscope, as described previously^14^. Experiments were performed in a flow chamber, placed on the microscope stage and illuminated with a collimated cold white LED (Thor Labs). The beads were visualized using 63X oil-immersion objective, attached with the nanofocusing piezo actuator (P-725, Physik instrumente). Images were processed through Ximea camera (MQ013MG-ON). Both the piezo positioning and data acquisition were measured by a multifunctional DAQ (Ni-USB-6289, National Instrument). Mechanical forces were applied through the generation of magnetic field by neodymium magnets, which are attached to the linear voice coil (Equipment Solutions) the force can be controlled by changing the position of the magnets. Detail information on force calibration, bead tracking and image processing were discussed previously by Chakraborty et al^17^. Experiments were performed in flow chamber with 1-10 nM protein in 1X PBS buffer (pH 7.2). Drugs were commercially purchased: methotrexate (Sigma-Aldrich).

### MD Simulation

Molecular dynamics simulations of protein with methotrexate (MTX, PubChem CID-126941) were performed using the NAMD 2.19 package using a concentration-dependent simulation protocol. The talin structure was taken from PDB ID 2L7A and the protonation states of the talin amino acid residues were determined by the H++ server. Four simulations were performed: one without MTX (control) and three with MTX concentrations: 1 μM, 40 μM, and 100 μM. The Automated Topology Builder (ATB) repository was used to generate force fields for the ligand. CHARMM 36 force field was used to study the protein-MTX interaction using the TIP3P water model. Initially, drug structures were stored randomly in a 10 x 10 x 10 nm cubic box. The systems were then solvated with ~32000, 31979, 31370, and 30796 TIP3P water molecules for control, 1, 40, and 100 μM MTX, respectively, in a 10 nm cubic box with 60 nm periodic boundary conditions. Na+ and Cl-ions were added to achieve neutrality and bring the final salt concentration to 150mM. The step size was kept fixed at 2 fs/time step. Energy minimization was performed using the sophisticated conjugate gradient and line search algorithm for 10000 steps (20 ps). Each system was equilibrated for 500,000 steps (1 ns). Temperature control of the system was performed using Langevin dynamics and a Berendsen pressure coupling was used for pressure control. The solvent density was maintained using a Generalized Born Implicit Solvent (GBIS) method with a coupling constant of 0.1 ps at 1 atm and 310 K to perform equilibration in NPT (number of atoms in the system, pressure of the system, and temperature of the systems). A cut-off of Coulomb forces was set with a switching function starting at 10 Å and reaching zero at 13 Å. We used full electrostatics with Particle-Mesh-Ewald (PME) summation to eliminate surface effects. Outputs of These equilibrated systems were then taken as inputs for constant velocity-steered molecular dynamics (SMD).

NAMD 2.19 was used to perform the SMD simulations with constant velocity pulling. Cα-atom of 781-aa residue was fixed, and an external force was applied to the Cα-atom of 911-aa residue (SMD-atom). A virtual harmonic spring attached to the SMD-atom, was pulled with a constant velocity v towards the direction of the vector linking the fixed and SMD-atom. The simulations were performed at 310 K using a constant velocity of 0.05 Å/ps. Simulations were run for 10 ns and end-to-end distance of the protein was checked for confirmation of complete unfolding. The force probed by the virtual spring was calculated using the equation: F = k (vt - (x0 - x)); where k = 7 kcal/mol/ Å^2^ (416 pN/Å), v is the velocity, t is time, and x and x0 are the 3D-vector defined position of the SMD-atom and the dummy atom at the opposite end of the spring, respectively. Each simulation was triplicated and the results from the simulations were visualized and analyzed in VMD.

Secondary structure perturbations, H-bonding, Salt-bridge interactions, and RMSD analysis of the simulations were performed using VMD Timeline plugin. The cut-off for H-bonding was set at 3.0 Å. The cut-off for short-range electrostatic interactions and long-range electrostatic interactions were chosen to be 5 Å and 10 Å respectively. Salt-bridge interactions were defined based on a cut-off distance < 3.5 Å.

### Cell lines and cell culture

HEK293T and HeLa cells were cultured in accord with the standard protocols in Dulbecco’s modified Eagle’s medium (DMEM, GIBCO) supplemented with 10% FBS (Himedia), and penicillin/streptomycin, (100 U/ml and 100 μg/ml, Invitrogen) at 37°C in a humidified incubator at 5% CO_2_. Trypsinization were performed when cells were grown to 80-90% confluency.

### Cell-shape measurement

Cells were seeded onto 24-well plates at 1 x 10^5^ per well. Methotrexate was administered to wells at 1 μM, 50 μM, and 100 μM conc. except the control well after overnight incubation at 37°C in a humidified 5% CO_2_ incubator. Images were taken using the bright-field microscopy at 10x (Evos, Thermofisher). Only the cells that were not clustered, were considered for analysis. cell shape index (circularity index) was determined for HEK293T and HeLa cultures by evaluating one frame from each *n* (no. of replicates, total of three images) per treatment for cell circularity measurement using ImageJ software. The shape index (circularity index) was calculated using the following formula:

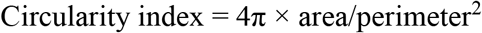

This is an established method to determine cell shape^43^. A complete circle would have a shape index of 1 whereas, a straight line would show an index of 0.

### Cell adhesion assay

Cells were treated with solvent only, 0.25 μM, 0.5 μM, 1 μM, 50 μM and 100 μM methotrexate (MTX). After 48 hours the cells were trypsinized, counted and used for the assay. A polystyrene-coated 96-well plate was pre-coated with 5 μg/ml fibronectin (Millipore). The working concentration (5 μg/ml) of fibronectin was prepared from the stock solution of 1 mg/ml. From the working solution of fibronectin, 200 μl per well was used for coating and the plates incubated overnight at 4°C temperature. All untreated and treated cells were seeded into these pre-coated 96-well plates, maintaining 1 x 10^5^ cells/well. The next day, the wells were blocked with 210 μL of 10 mg/mL BSA for 30 minutes at room temperature (BSA was heat denatured at 70°C for 30 minutes). Untreated control cells were also blocked using the same procedure. Then the blocked cells were incubated for 30 minutes at 37°C in a CO_2_ incubator. The cells in each well were washed three times with PBS to remove cell debris. Then the cells were fixed with 4% PFA (paraformaldehyde) for 15-20 minutes, followed by washing with PBS three times. 100 μl of 0.1% crystal violet was added to each well and incubated for 20 minutes at room temperature. 10% GAA (glacial acetic acid) was added to each well after washing three times with PBS, and the OD of each well was measured at 595 nm in a multiplate UV/vis spectrometer (CLARIOstar). Data are presented as mean S.E.M; (n=3).

### In-vitro scratch-induced migration assay

HEK293T and HeLa cells were taken out of culture in their 6^th^ passage upon reaching ~90% confluency. The cells were seeded at 2×10^5^ per well in 12-well plates and grown to 95-100% confluence. A linear scratch wound was made by a sterile p200 tip in the adherent confluent monolayer of the cells. Wells were washed with sterile 1X PBS four times to remove any cell debris. Cells were treated with only solvent, 0.25 μM, 0.5 μM, 1 μM, 50 μM, and 100 μM of Methotrexate (MTX). The first imaging was done immediately after drug administration (0 hr). The treated cells were kept at 37°C in a humidified incubator at 5% CO_2_ for 24 hrs for the second imaging. The closure extent of the cell-free wounded area was photographed by bright-field microscopy at 10x (Evos, Thermofisher). Wound closure was quantified by Fiji ImageJ software. The wound closure percentage were calculated as:

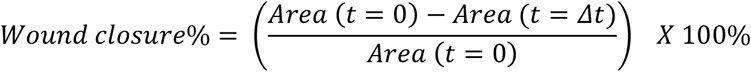

t=0 represents the first imaging time and t= Δ*t* represents the final imaging time. The areas of wounds are represented in pixel^2^. We considered comparing migration rates of both cells in different conc. or in absence of MTX. Thus, the migration rate was calculated as follows:

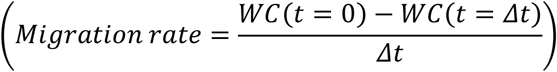

where, WC = wound closure percentage and Δt = time of experiment (migration time). Data are represented as means ±S.E.M; (n=3).

### Plasmids and antibodies

| Name | Obtained from | Cat. No. |
| --- | --- | --- |
| Talin-EGFP plasmid construct (mammalian) | Kind gift from Dr. Michael Davidson |  |
| Vinculin-mCherry plasmid construct (mammalian) | Kind gift from Dr. Michael Davidson |  |
| LATS1 rabbit monoclonal primary antibody | Cell signaling Technology | 3477 |
| Mst1 rabbit monoclonal primary antibody | Cell signaling Technology | 3682 |
| Yap1 rabbit monoclonal primary antibody | Cell signaling Technology | 14074 |
| pYAP rabbit monoclonal primary antibody | Cell signaling Technology | 13008 |
| β-actin mouse primary antibody | Santa Cruz Biotechnology | Sc-47778 |
| HRP-conjugated anti-rabbit secondary antibody (IgG) | Cell signaling Technology | 7074 |
| HRP-conjugated anti-mouse secondary antibody (IgG) | Cell signaling Technology | 7076 |

### Co-transfection of Talin and Vinculin constructs

HEK293T and HeLa cells were seeded on microscopic coverslips in a 12 well plate (1.5 × 10^5^ cells/well). Cells were co-transfected with mCherry-Vinculin and EGFP-Talin1 (500ng each, plasmids were kind gift from Dr. Michael Davidson), using Lipofectamine 3000 transfection reagent (Invitrogen) with opti-MEM (transfection medium), as per manufacturer’sinstructions. After 24 h of transfection, the transfection medium was replaced with a complete medium followed by scratch wound on the monolayer as previously described. Treatment was introduced as administration of only solvent, 1, 40, and 100 μM of MTX to the corresponding wells and cells were incubated for another 12 h.

### Confocal imaging

For fluorescence confocal imaging, cells were washed once with PBS to remove residual media and fixed using 4% paraformaldehyde (PFA) in PBS for 15 minutes. The nuclei were stained using Hoechst 33342 dye (Invitrogen) at 1:2000 dilution in PBS for 5-10 minutes at room temperature. The cells were washed twice with PBS and mounted in Fluoromount-G mounting medium (Invitrogen) for imaging, as per the manufacturer’s instructions. Only the areas near the scratches were captured to only consider the cells with intrinsically generated traction due to migration. Fluorescent images were captured with a confocal laser microscope (Leica TCS SP8) and analyzed using the Coloc2 plugin of Fiji ImageJ software (https://imagej.net/plugins/coloc-2). As we have used fluorescence tagged plasmid overexpression system, we wanted to only cover the focal adhesion or adjacent regions. Therefore, in each image, cytoplasmic regions were omitted using selection tool of ImageJ.

Data are represented as means ±S.E.M; n=3. Focal adhesion measurements were performed according to the protocol given by Horzum et al^44^.

### Western blot analysis

Harvested cells were taken out of the −80° C temperature and thawed. They were given a quick spin, and the PBS was discarded from each tube and again placed in the ice. 1 ml lysis mix was prepared by adding 900 μl RIPA lysis buffer (150 mM NaCl, 1% NP-40, 0.5% sodium deoxycholate, 0.1% SDS, and 10 mM Tris–HCl pH 8.0), 100 μl 10X PPIC (Phosphatase and Protease Inhibitor Complex), and 1 mM PMSF. The lysis mix was added to each tube and kept in ice for 30 minutes with occasional vortex after every 10 minutes. Then, the samples were centrifuged at 12000g for 10 minutes at 4°C. The supernatants were taken, and the protein concentrations in each tube were measured by Bradford assay using Quick Start Bradford protein assay kit (Bio-Rad). An equal amount of each sample (30-35 μl), 4-5 μl Page Ruler Pre-stained Protein Ladder (Thermo Scientific), and 10 μl of 4X Dye (last three grooves) were loaded in SDS PAGE. SDS-PAGE was performed using Bio-Rad Mini-PROTEAN Tetra Cell. After SDS-PAGE, proteins were transferred to Immun-Blot PVDF (polyvinylidene fluoride) membranes (Bio-Rad) with transfer buffer by semi-wet electro transfer using trans-Blot Turbo transfer system (Bio-Rad). Blocking was done using a blocking buffer before primary antibody incubation. Antibodies to the following proteins were used: pYAP, Yap/TAZ, LATS1, β-Actin, and MST. β-actin was used as the loading control. For primary antibody, 1:1000 dilution and secondary antibody, 1:5000 dilution was used in each case. For each sample, after primary antibody addition, the sample was incubated overnight at 4°C. The membrane was washed with PBS-T (Tween-20, Merck), three times every 10 minutes and incubated for 2 hours with the HRP-conjugated secondary antibodies. After incubation, the membrane was washed with PBS-T, three times, 10 minutes each. The color was developed in the membrane using a solution of 1:1 luminol: peroxide in a dark room. Chemiluminescent images were taken in the ChemiDoc MP imaging system (Bio-Rad).

Quantifications were performed using ImageJ imaging software and graphs were generated using Origin Pro. For western blot protein expression quantitation, we preferred normalization of the signal from protein of interest with that of reference protein (β-actin). To determine the normalization factor for each lane that was used to normalize the experimental area values for the proteins of interest, the highest signal detected for the housekeeping protein (β-actin) was identified. The value of this band was used to normalize the rest of the β-actin bands on the blot. To determine the lane normalization factor (LNF) we employed the formula:

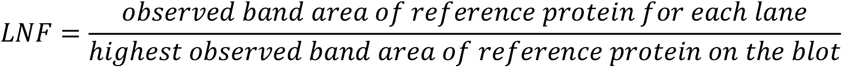

To calculate the normalized signal of each experimental target band, the observed signal intensities of each experimental target band should be divided by the lane normalization factor of the corresponding lane of that particular target band:

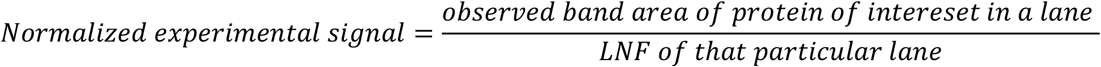

The experiments were triplicated and results were shown as Mean ± S.E.M.

### Genomic analysis

For cell-line specific genomic experiments, the IC50 of 481 drugs in 1001 cell lines and their corresponding mRNA gene expression from the CTRP (Cancer Therapeutics Response Portal), and the IC50 of 265 drugs in 860 cell lines and the respective mRNA expression from GDSC (Genomics of Drug Sensitivity in Cancer) were collected and merged. The mRNA expression data and the drug sensitivity data were merged to obtain sensitivity-specific mRNA expression values. Only the data for methotrexate treated samples where MTX conc. was kept within 1 μM and 50 μM, were selected. Pearson correlation coefficient was measured to obtain the correlation between gene mRNA expression and drug IC50. The P-value was adjusted by FDR and considered significant only if FDR<0.05. For mRNA expression, major genes of Talin-YAP Hippo signaling and their transcriptomic products were considered.

For real world genomic data, we downloaded TCGA genomics data and the corresponding treatment data from cBioportal. Only sample IDs which were reported to be treated with methotrexate, were considered (n=48). We analyzed expression of the major genes of Talin-YAP Hippo signaling and their transcriptomic products using their z-score value. For comparative analysis, both the tumor and normal data were considered. The p-value was adjusted by FDR and considered significant only if FDR<0.05.

### Statistics and data analysis

At least three independent experiments were performed in each analysis. Samples were randomly selected as control and treatment groups in the studies involving drug molecules. Sample sizes in genomic analysis were confirmed using power analysis (considered if power >80%). Comparative analyses were performed using either one-way ANOVA with Tukey’s multiple comparison and Bonferroni Post Hoc test or student t-test. Two sample t-tests were performed to compare two groups with same number of data whereas paired t-test was performed for comparing two groups with unequal amount of data. In each case a p-value < 0.05 was considered significant. To find the correlation between two events Pearson’s correlation coefficient was calculated. Gene expression values were converted to z-score for analysis and z-score ± 1.96 was considered significant. FDR > 0.05 was considered significant to find correlation. All the p-values and number of samples are mentioned in the figures or figure legends if not specified otherwise. Simulations were performed and analyzed in NMRbox. The single molecule and simulation data acquisition and analysis were performed with Igor Pro 8.0 software (Wavemetrics). Sequencing data from the databases were analyzed and plotted using different libraries of python 3.0. Simulation results are visualized, and analysed with VMD, PyMOL, and Biovia Discovery studio visualizer. Cell-based analysis was performed using Fiji ImageJ software. The fluorescence images were processed using Leica Software LAS-X 3.7.6 and analyzed using ImageJ.

## Data availability

Cell line specific drug-treatment and corresponding genomic data are available from GDSC (https://www.cancerrxgene.org/) and CTRP (https://portals.broadinstitute.org/ctrp.v2.1/). The TCGA treatment datasets can be obtained from GDC (https://portal.gdc.cancer.gov/) or cBioPortal (https://www.cbioportal.org/datasets). All other data produced during this study are available from the corresponding author upon reasonable request.

## Author contribution

S.H., D.C. designed the project. M.B., D.C., S.D., D.M., K.K.A. performed the experiments. M.B., D.C., S.D., D.M., S.P. analyzed the data. D.C., S.H., and S.C. wrote the manuscript.

## Acknowledgments

We thank Ashoka University, Trivedi School of Biosciences and Mphasis foundation for support and funding. D.C. thanks Department of Biotechnology, India for DBT JRF. S.H. thanks DBT Ramalingaswami Fellowship and DST SERB Core Research Grant for funding. We would like to thank IIT Delhi, CRF for confocal facility.

## Reference

1. Chen, A. R., Wang, Y. M., Lin, M. & Kuo, D. J. High-Dose Methotrexate in Pediatric Acute Lymphoblastic Leukemia: Predictors of Delayed Clearance and the Effect of Increased Hydration Rate on Methotrexate Clearance. Cureus (2020) doi:10.7759/cureus.8674.

2. Inaba, H. et al. Clinical and radiological characteristics of methotrexate-induced acute encephalopathy in pediatric patients with cancer. Annals of Oncology 19, 178–184 (2008).

3. Wojtuszkiewicz, A. et al. Folylpolyglutamate synthetase splicing alterations in acute lymphoblastic leukemia are provoked by methotrexate and other chemotherapeutics and mediate chemoresistance: Folylpolyglutamate Synthetase Splicing Alterations in ALL. Int. J. Cancer 138, 1645–1656 (2016).

4. Polesie, S. et al. Use of methotrexate and risk of skin cancer: a nationwide case–control study. Br J Cancer 128, 1311–1319 (2023).

5. Kreher, M. A., Konda, S., Noland, M. M. B., Longo, M. I. & Valdes-Rodriguez, R. Risk of melanoma and nonmelanoma skin cancer with immunosuppressants, part II: Methotrexate, alkylating agents, biologics, and small molecule inhibitors. Journal of the American Academy of Dermatology 88, 534–542 (2023).

6. Díaz-Valdivia, N. I. et al. Anti-neoplastic drugs increase caveolin-1-dependent migration, invasion and metastasis of cancer cells. Oncotarget 8, 111943–111965 (2017).

7. Goult, B. T., Yan, J. & Schwartz, M. A. Talin as a mechanosensitive signaling hub. J Cell Biol 217, 3776–3784 (2018).

8. Yao, M. et al. The mechanical response of talin. Nature Communications 7, 11966 (2016).

9. Sun, Z., Guo, S. S. & Fässler, R. Integrin-mediated mechanotransduction. J Cell Biol 215, 445–456 (2016).

10. Tapia-Rojo, R., Alonso-Caballero, Á. & Fernández, J. M. Talin folding as the tuning fork of cellular mechanotransduction. PNAS 117, 21346–21353 (2020).

11. Tapia-Rojo, R., Alonso-Caballero, A. & Fernandez, J. M. Direct observation of a coil-to-helix contraction triggered by vinculin binding to talin. Science Advances 6, eaaz4707 (2020).

12. Atherton, P. et al. Vinculin controls talin engagement with the actomyosin machinery. Nat Commun 6, 10038 (2015).

13. Yao, M. et al. Mechanical activation of vinculin binding to talin locks talin in an unfolded conformation. Scientific Reports 4, 4610 (2014).

14. Popa, I. et al. A HaloTag Anchored Ruler for Week-Long Studies of Protein Dynamics. J. Am. Chem. Soc. 138, 10546–10553 (2016).

15. Valle-Orero, J. et al. Mechanical Deformation Accelerates Protein Ageing. Angew Chem Int Ed Engl 56, 9741–9746 (2017).

16. Chakraborty, S. et al. Connecting conformational stiffness of the protein with energy landscape by a single experiment. Nanoscale 14, 7659–7673 (2022).

17. Chakraborty, S., Chaudhuri, D., Banerjee, S., Bhatt, M. & Haldar, S. Direct observation of chaperone-modulated talin mechanics with single-molecule resolution. Commun Biol 5, 1–14 (2022).

18. Chaudhuri, D., Banerjee, S., Chakraborty, S., Chowdhury, D. & Haldar, S. Direct Observation of the Mechanical Role of Bacterial Chaperones in Protein Folding. Biomacromolecules 23, 2951–2967 (2022).

19. Haldar, S., Tapia-Rojo, R., Eckels, E. C., Valle-Orero, J. & Fernandez, J. M. Trigger factor chaperone acts as a mechanical foldase. Nature Communications 8, 668 (2017).

20. Haining, A. W. M., von Essen, M., Attwood, S. J., Hytönen, V. P. & del Río Hernández, A. All Subdomains of the Talin Rod Are Mechanically Vulnerable and May Contribute To Cellular Mechanosensing. ACS Nano 10, 6648–6658 (2016).

21. Stannard, A. et al. Molecular Fluctuations as a Ruler of Force-Induced Protein Conformations. Nano Lett. 21, 2953–2961 (2021).

22. Gingras, A. R. et al. Mapping and Consensus Sequence Identification for Multiple Vinculin Binding Sites within the Talin Rod. Journal of Biological Chemistry 280, 37217–37224 (2005).

23. Rahikainen, R. et al. Mechanical stability of talin rod controls cell migration and substrate sensing. Sci Rep 7, 3571 (2017).

24. Lemke, S. B., Weidemann, T., Cost, A.-L., Grashoff, C. & Schnorrer, F. A small proportion of Talin molecules transmit forces at developing muscle attachments in vivo. PLOS Biology 17, e3000057 (2019).

25. Liu, J. et al. Talin determines the nanoscale architecture of focal adhesions. Proceedings of the National Academy of Sciences 112, E4864–E4873 (2015).

26. Beaty, B. T. et al. Talin regulates moesin-NHE-1 recruitment to invadopodia and promotes mammary tumor metastasis. J Cell Biol 205, 737–751 (2014).

27. Goult, B. T. et al. RIAM and Vinculin Binding to Talin Are Mutually Exclusive and Regulate Adhesion Assembly and Turnover *. Journal of Biological Chemistry 288, 8238–8249 (2013).

28. Campanale, J. P., Sun, T. Y. & Montell, D. J. Development and dynamics of cell polarity at a glance. Journal of Cell Science 130, 1201–1207 (2017).

29. Pasqualato, A. et al. Shape in migration: Quantitative image analysis of migrating chemoresistant HCT-8 colon cancer cells. Cell Adhesion & Migration 7, 450–459 (2013).

30. Janiszewska, M., Primi, M. C. & Izard, T. Cell adhesion in cancer: Beyond the migration of single cells. J Biol Chem 295, 2495–2505 (2020).

31. Haining, A. W. M. et al. Mechanotransduction in talin through the interaction of the R8 domain with DLC1. PLOS Biology 16, e2005599 (2018).

32. Sun, Z., Costell, M. & Fässler, R. Integrin activation by talin, kindlin and mechanical forces. Nat Cell Biol 21, 25–31 (2019).

33. Kim, D. & Wirtz, D. Focal adhesion size uniquely predicts cell migration. FASEB j. 27, 1351–1361 (2013).

34. Elosegui-Artola, A. et al. Mechanical regulation of a molecular clutch defines force transmission and transduction in response to matrix rigidity. Nat Cell Biol 18, 540–548 (2016).

35. Mason, D. E. et al. YAP and TAZ limit cytoskeletal and focal adhesion maturation to enable persistent cell motility. Journal of Cell Biology 218, 1369–1389 (2019).

36. Jang, J.-W. et al. RAC-LATS1/2 signaling regulates YAP activity by switching between the YAP-binding partners TEAD4 and RUNX3. Oncogene 36, 999–1011 (2017).

37. Yang, W. et al. Genomics of Drug Sensitivity in Cancer (GDSC): a resource for therapeutic biomarker discovery in cancer cells. Nucleic Acids Research 41, D955–D961 (2012).

38. Basu, A. et al. An Interactive Resource to Identify Cancer Genetic and Lineage Dependencies Targeted by Small Molecules. Cell 154, 1151–1161 (2013).

39. Kim, M.-K., Jang, J.-W. & Bae, S.-C. DNA binding partners of YAP/TAZ. BMB Rep. 51, 126–133 (2018).

40. Edwards-Hicks, J. et al. MYC sensitises cells to apoptosis by driving energetic demand. Nat Commun 13, 4674 (2022).

41. Atherton, P. et al. Relief of talin autoinhibition triggers a force-independent association with vinculin. Journal of Cell Biology 219, e201903134 (2020).

42. Rahikainen, R., Öhman, T., Turkki, P., Varjosalo, M. & Hytönen, V. P. Talin-mediated force transmission and talin rod domain unfolding independently regulate adhesion signaling. Journal of Cell Science jcs.226514 (2019) doi:10.1242/jcs.226514.

43. Schutte, S. C. & Taylor, R. N. A tissue-engineered human endometrial stroma that responds to cues for secretory differentiation, decidualization, and menstruation. Fertility and Sterility 97, 997–1003 (2012).

44. Horzum, U., Ozdil, B. & Pesen-Okvur, D. Step-by-step quantitative analysis of focal adhesions. MethodsX 1, 56–59 (2014).

